# Whole-Cell Proteomics Identifies Novel Regulators of Ciliogenesis Beyond the Axoneme

**DOI:** 10.1101/2025.05.20.655211

**Authors:** Xiaolu Xu, Yanbao Yu, Tony Zheng, Fiona Clark, Jean Ross, Neha Sindhu, Andre L P Tavares, John B Wallingford, Shuo Wei, Jian Sun

## Abstract

Motile cilia generate directional fluid flow critical for development and tissue homeostasis, and their dysfunction underlies a range of human diseases. Prior proteomic studies have focused on isolated axonemes, leaving the regulatory components of ciliogenesis located in other cellular compartments of multiciliated cells (MCCs) largely unidentified. To bridge this gap, we performed whole-cell proteomic profiling of MCCs in *Xenopus* using an inducible system that enables selective MCC-enrichment. Leveraging a single-vessel in-cell proteomics approach with ultra-high sample preservation, we identified over 9,000 proteins, including 832 upregulated in MCCs. These encompassed both established ciliary components and novel candidates localized to distinct ciliary structures. Functional validation confirmed the involvement of previously uncharacterized proteins in ciliogenesis. This study presents a high-resolution, whole-cell proteome of vertebrate MCCs, providing critical insights into the molecular networks that govern ciliogenesis and offering a valuable resource for understanding the mechanisms of cilia-related disorders.

## INTRODUCTION

Motile cilia are microtubule-based organelles that generate directional fluid flow, playing fundamental roles in biological processes such as mucus clearance in the respiratory tract, cerebrospinal fluid circulation, and embryonic development (Brooks and Wallingford, 2014; Boutin and Kodjabachian, 2019; Fliegauf et al., 2007; Spassky and Meunier, 2017). Dysfunction of motile cilia results in ciliopathies, a group of disorders associated with chronic respiratory infections, hydrocephalus, heterotaxy, and infertility (Nigg and Raff, 2009; Reiter and Leroux, 2017). Despite significant progress in understanding ciliary structures and functions, many molecular regulators of ciliogenesis remain unidentified.

Proteomic approaches have been instrumental in mapping the molecular composition of cilia, leading to the identification of numerous proteins involved in cilia assembly and function (Li et al., 2004; Chen et al., 2023). Early proteomic analyses of ciliary axonemes from human bronchial epithelial cells identified over 200 proteins, including core ciliary components such as tubulins and dyneins (Ostrowski et al., 2002). Subsequent studies expanded the ciliary proteome, identifying 352 axonemal proteins, including multiple tubulin isoforms and various non-tubulin constituents (Blackburn et al., 2017). In *Xenopus laevis*, proteomic characterization of isolated cilia has identified many putative ciliary proteins, including keratin 17, which plays a crucial role in ciliogenesis (Sim et al., 2020). Comparative studies in *Chlamydomonas reinhardtii* and *Trypanosoma* have further revealed evolutionarily conserved ciliary proteins, identifying homologs of vertebrate disease-related proteins such as Polycystin 2, Fibrocystin, and Hydin (Boesger et al., 2009; Broadhead et al., 2006; Pazour et al., 2005; Subota et al., 2014).

While these proteomic studies using isolated cilia have provided valuable insights into the axonemal composition, they focus exclusively on the ciliary axoneme while overlooking other critical cellular components involved in ciliogenesis. Multiciliated cell (MCC) function is regulated not only by axonemal proteins but also by basal bodies, cytoplasmic factors, and nuclear components (Brooks and Wallingford, 2014; Spassky and Meunier, 2017). The reliance on deciliation, axoneme isolation, or shedding in previous studies has largely overlooked these regulatory elements. To address this limitation, a more comprehensive proteomic approach is desired to capture the entire molecular network governing cilia formation and function. Thus, expanding proteomic profiling to whole MCCs is essential for providing a broader perspective on ciliogenesis by incorporating proteins from all cellular compartments involved in MCCs development.

We recently developed On-Filter In-Cell (OFIC)-based sample preparation as a robust strategy for high-sensitivity proteomic analysis, reducing processing variability and enabling efficient detection of low-abundance proteins, particularly in complex biological samples including *Xenopus* embryos (Martin et al., 2024; Elsayyid et al., 2025; Sun et al., 2025). Unlike conventional lysis-based approaches, which require multiple processing steps that may introduce sample loss and biases, OFIC preserves the spatial organization of proteins while maintaining biochemical integrity. This method drastically streamlines proteomic workflows, making it particularly advantageous for low sample inputs and enabling more comprehensive investigations of MCCs at a systems level.

By leveraging *Xenopus* animal cap–derived mucociliary organoids in combination with OFIC sample processing and mass spectrometry, we generated a high-resolution, cell type–enriched proteomic profile of whole MCCs. Functional validation through in situ hybridization, immunostaining, and gene knockdown revealed several previously uncharacterized proteins that are essential for MCC maintenance and ciliogenesis. This comprehensive characterization of the MCC proteome provides valuable insights into the molecular networks driving ciliogenesis. Our findings not only broaden the catalogs of known ciliary proteins but also identify novel regulatory factors, offering new perspectives for understanding cilia-related diseases and advancing potential therapeutic approaches.

## RESULTS

### Generation of a Comprehensive MCC Proteome Using *Xenopus* Animal Cap Organoids

The embryonic epidermis of *Xenopus* has emerged as a powerful vertebrate model for studying normal ciliogenesis and ciliopathies as its cellular composition, which includes both multiciliated and secretory cells, closely resembling the mammalian airway epithelium (Walentek and Quigley, 2017; Walentek, 2021; Lee et al., 2023). Many key proteins involved in mucociliary differentiation and function are conserved between *Xenopus* and mammals (Werner and Mitchell, 2012, 2013). Additionally, *Xenopus* animal caps, pluripotent cell sheets derived from the animal pole of blastula-stage embryos, serve as an effective ex vivo model. When cultured as explants, these organoids undergo default mucociliary differentiation, faithfully recapitulating early developmental events and providing a tractable system for investigating MCC biology (Angerilli et al., 2018; Lee et al., 2023).

A major obstacle in whole-cell proteomic studies of MCCs is the challenge of distinguishing MCC-specific proteins from the other cellular proteins that can obscure the identification of potential regulators of ciliogenesis. To overcome this obstacle, we employed *Xenopus laevis* animal cap organoids as an *ex vivo* system to model MCC differentiation. Through ectopic expression of MCIDAS, a master regulator of centriole assembly and ciliogenesis (Ma et al., 2014; Stubbs et al., 2012), we robustly induced MCC cell fate in animal caps and enriched ciliary proteins. To selectively enhance MCC differentiation, MCIDAS was fused with glucocorticoid receptor (GR) and activated upon treatment with dexamethasone (Dex). This strategy created an inducible MCC gain-of-function model, in which Dex treatment robustly induced MCC formation within the organoids. By comparing the proteomic profiles of Dex-treated and untreated animal cap organoids, we sought to enrich for proteins upregulated in MCCs, thereby reducing the background noise and enhancing the specificity of the proteomic dataset.

MCIDAS-GR was overexpressed at one-cell stage, followed by dissection of animal caps at blastula stage (stage 9; Figure 1A). To minimize early developmental toxicity, Dex was not added to the cultured explants until stage 11 to activate MCIDAS-GR, which resulted in robust MCC differentiation by stage 26 (Figure 1A’). Compared to untreated controls, Dex-treated organoids exhibited a marked increase in MCC formation, as confirmed by acetylated tubulin staining (Figure 1B; Figure S1A).

**Figure 1.**
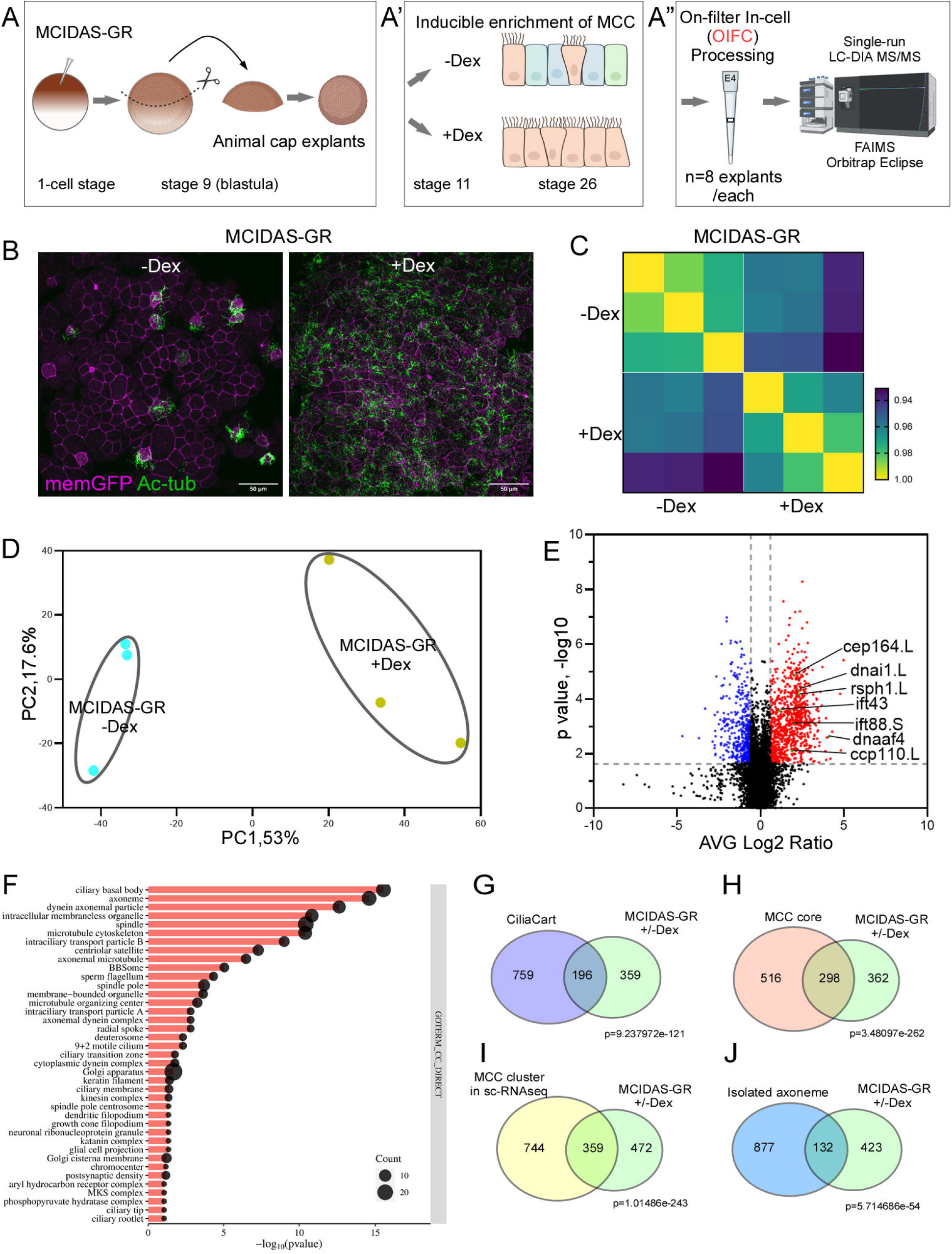
Profiling MCC proteome using *Xenopus* animal cap organoids with OFIC approach. (A-A”) Illustrative workflow of the proteomic analysis of *X. laevis* animal caps enriched with MCCs. Upon fertilization, embryos were injected with mRNA encoding MCIDAS-hGR and then cultured to stage 9, when animal caps were dissected and cultured in vitro to generate organoids. With addition of dexamethasone, the majority of epidermal cells undergo differentiation toward a MCC fate, as compared to untreated animal caps. The organoids were cultured to stage 26 and were fixed with methanol. E4tip was used for OFIC processing. The mass spectrometric analysis was performed using Orbitrap Eclipse. (B) Organoids were fixed at stage 26 and stained for acetylated tubulin (Ac-tub). mRNA encoding a membrane-bound form of GFP (memGFP) was co-injected as a tracer to label targeted cells. Representative confocal images of the epidermis for each condition were shown. Scale bar: 10 μm. (C) Pearson correlation analysis. (D) Principal component analysis. (E) Volcano plot of comparison between Dex-treated organoids and untreated ones. Well-characterized ciliogenesis-related proteins were labeled. (F) Gene ontology cellular component analysis of significantly upregulated proteins in (E). (G) Venn diagram showing the overlap between significantly upregulated proteins in (E) and CiliaCart database. Note that protein counts were adjusted to reflect human orthologs mapped from our dataset. The p-value was calculated using a hypergeometric distribution test. (H) Venn diagram showing the overlap between significantly upregulated proteins in (E) and the MCC core gene database. Note that protein counts from our dataset were adjusted by merging L and S isoforms of the same gene and excluding uncharacterized proteins. The p-value was calculated using a hypergeometric distribution test. (I) Venn diagram showing the overlap between significantly upregulated proteins in (E) and the MCC sc-RNAseq database. The p-value was calculated using a hypergeometric distribution test. (J) Venn diagram showing the overlap between significantly upregulated proteins in (E) and the axoneme proteome. Note that protein counts were adjusted to reflect human orthologs. The p-value was calculated using a hypergeometric distribution test.

### High-Depth Proteomic Profiling of MCCs Using OFIC Processing and Mass Spectrometry

Given the small size of animal cap organoids (diameter <0.5 mm), conventional lysis-based proteomic workflows require a large number of samples and multiple processing steps, potentially leading to inefficiencies and protein loss. To overcome these limitations, OFIC processing was used, enabling direct protein digestion in fixed cells while minimizing sample loss and enhancing proteomic depth (Elsayyid et al., 2025; Martin et al., 2024; Sun et al., 2025). Unlike conventional methods, which require nearly 100 organoids per replicate, the OFIC-based approach enabled robust proteomic analysis using only eight organoids per replicate across three biological replicates (Figure 1A”).

Liquid chromatography (LC)-MS analysis identified over 9,000 proteins, indicating high proteome coverage. The Pearson correlation averaged from 0.97 to 0.98 within three biological replicates (Figure 1C), confirming high reproducibility of the OFIC method. Principal Component Analysis (PCA) showed distinct separation of Dex+ from Dex− groups (Figure 1D), suggesting substantial proteomic differences between MCC-enriched and control conditions. We identified 832 upregulated proteins in Dex+ organoids compared to Dex-controls, including well-known regulators of ciliogenesis (e.g., CP110, CEP164) and ciliary motility (e.g., RSPH1, DNAI1, DNAAF4) (Figure 1E) (Guo et al., 2022; Kott et al., 2013a; Schmidt et al., 2012; Spektor et al., 2007; Zariwala et al., 2006). As expected, Dex treatment alone had minimal effects on the organoid proteome in the absence of MCIDAS induction (Figure S1B and S1C).

Our MCC proteomic dataset was significantly enriched for cilia-associated GO terms (Figure 1F). To further assess dataset quality, we compared our MCIDAS-GR-upregulated protein set with four established cilia/MCC reference datasets: CiliaCarta, the MCC Core Gene list, an MCC single-cell RNA-seq dataset, and Rfx2 direct target genes (Van Dam et al., 2019; Quigley and Kintner, 2017; Lee et al., 2023; Chung et al., 2014). We identified highly significant overlap with each of these datasets (Figure 1G-I; Figure S1D), supporting the biological relevance of our MCC proteome.

We also benchmarked our dataset against a published *X. laevis* axoneme-enriched proteome (Sim et al., 2020). In this comparison, 132 proteins overlapped with the axoneme dataset, whereas 432 MCC-upregulated proteins were not detected in the axoneme preparation (Figure 1J). These non-overlapping proteins likely represent MCC-associated factors beyond the axoneme, including basal body-, cytoplasmic-, and nuclear-enriched components, underscoring the ability of whole-cell profiling to reveal regulatory proteins missed by axoneme-focused proteomics.

### Localization of Novel Candidate Proteins to Diverse Ciliary Structures

To validate the novel candidates, three uncharacterized proteins—Coiled-Coil Domain-Containing 30 (Ccdc30), Membrane Occupation and Recognition Nexus-motif protein 2 (Morn2), WD Repeat Domain 88 (Wdr88) were selected for further investigation using *Xenopus tropicalis*, a species closely related to *X. laevis*. Unlike *X. laevis*, which is an allotetraploid, *X. tropicalis* has a diploid genome and is more convenient for loss-of-function studies (Hellsten et al., 2010; Session et al., 2016). Whole-mount in situ hybridization confirmed that *ccdc30, morn2*, and *wdr88* are specifically expressed in epidermal MCCs of *X. tropicalis* embryos, as indicated by co-localization with acetylated tubulin (Figure 2A-C).

**Figure 2.**
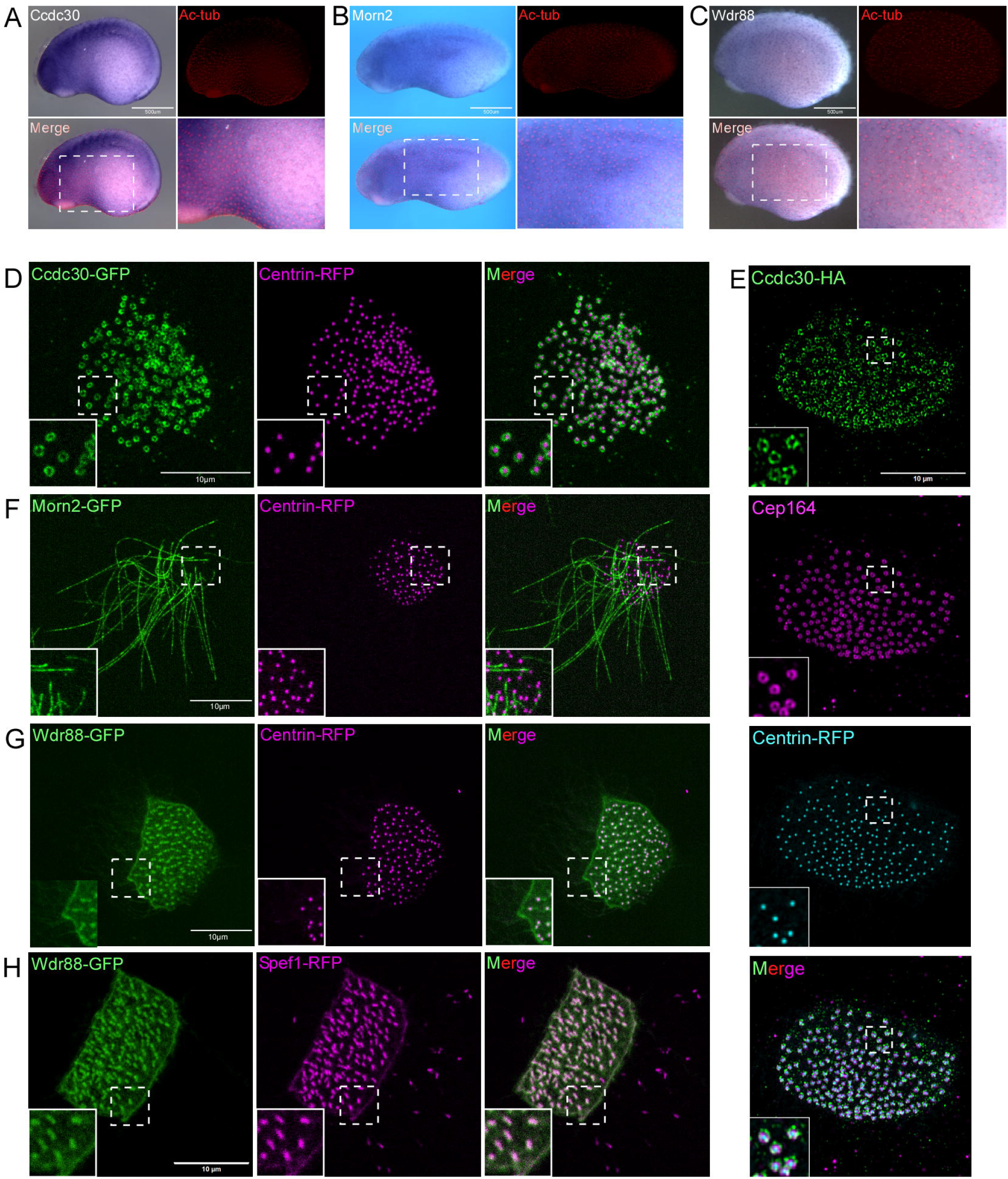
Expression and subcellular localization of candidate novel cilia-related proteins in epidermal MCCs. (A-C) Whole-mount in situ hybridization showing the expression of *ccdc30, morn2* and *wdr88* in epidermal MCCs at stage 22-24, followed by immunostaining with anti– acetylated tubulin to mark motile cilia. (D,F,G) Live imaging of MCCs overexpressing GFP-tagged candidate proteins with Centrin4-RFP marking basal bodies. Ccdc30-GFP (D) formed a ring-like structure surrounding basal bodies, resembling basal body appendages. Morn2-GFP (F) localized along the axoneme but was absent from basal bodies. Wdr88-GFP (G) was associated with basal bodies and exhibited an expanded appendage-like pattern overlapping with Centrin4-RFP. (E) Immunostaining of MCCs for Ccdc30-HA and the distal appendage marker Cep164 shows that Ccdc30 forms a ring-like structure with limited spatial overlap with Cep164. Scale bar: 10 μm. (H) Wdr88-GFP co-localized with rootlet marker Spef1-RFP. Scale bar: 10 μm.

We then assessed the subcellular localization using GFP-tagged versions of each candidate in MCCs. Ccdc30 is localized as a striking ring-like structure surrounding the centrioles, resembling basal body appendages (Figure 2D). We then assessed its precise positioning to centriole appendages by co-staining with the distal appendage marker protein Cep164 (Schmidt et al., 2012). Ccdc30 showed minimal overlap with Cep164 and appeared as a larger, more distally positioned ring structure as compared to Cep164 (Figure 2E and Video S1), highlighting its potential role in basal body organization and appendage assembly. Live imaging of both proteins yielded the same pattern (Video S2). By comparison, Morn2 was detected along the axoneme but excluded from basal bodies (Figure 2F). Wdr88 localized to basal body–associated structures (Figure 2G) and co-localized with Spef1/Clamp (Figure 2H), a marker of ciliary rootlet (Park et al., 2008).

### Morn2 is a Novel Regulator of Ciliogenesis

Given that Morn-domain containing proteins have been implicated in ciliopathies and localize to ciliary structures (Leung et al., 2025; Onoufriadis et al., 2014; Kott et al., 2013b), we selected Morn2 for further functional analysis. To assess ciliary function, we performed embryo gliding assays. Coordinated beating of epidermal MCCs generates directional fluid flow that propels the embryo, and the speed of this movement serves as an indicator of proper motile ciliary function. Knockdown of Morn2 by a translation-blocking antisense morpholino oligonucleotide (MO) significantly impaired embryo gliding, which was rescued by co-injection of Morn2 MO-resistant mRNA (Figure 3A and 3A’), indicating an essential role for this protein in MCC function.

**Figure 3.**
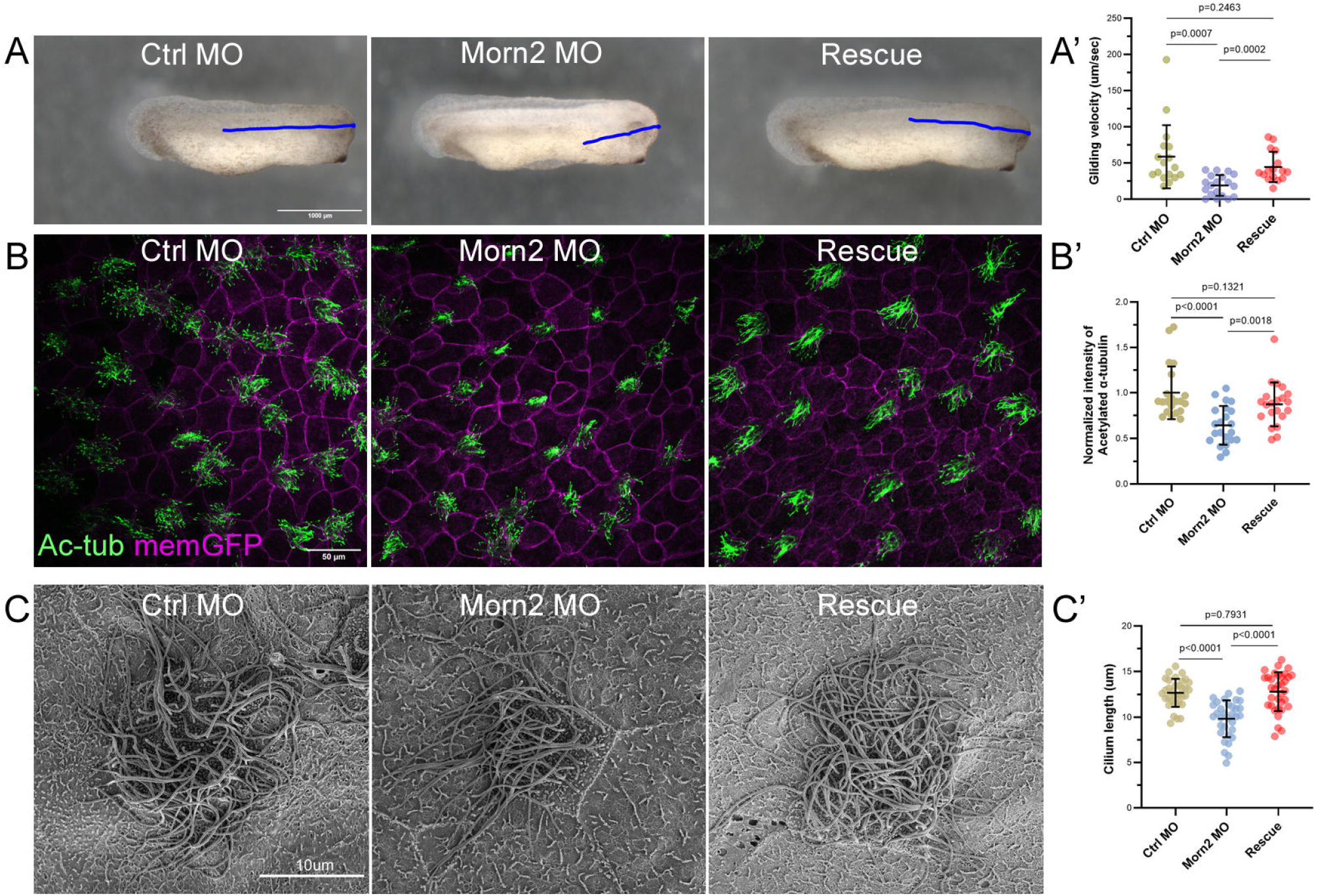
Morn2 is essential for ciliogenesis of MCCs. (A) Gliding assay showing that Morn2 knockdown significantly impaired embryo motility, a phenotype rescued by co-injection of Morn2 MO-resistant mRNA. Scale bar: 1000 μm. (A’) Quantification of gliding velocity in A from three independent experiments with unpaired two-tailed t-tests is shown in B. Mean ± s.d. values are presented. ns: no statistical differences between the groups. (B) Immunostaining for acetylated tubulin reveals a dramatic loss of motile cilia in Morn2 morphants, which is rescued by Morn2 mRNA re-expression. Scale bar: 50 μm. (B’) Quantification of normalized intensity of acetylated-tubulin in C from three independent experiments with unpaired two-tailed t-tests is shown in D. Mean ± s.d. values are presented. (C) Scanning electron microscopy (SEM) of MCCs shows a marked reduction in cilia number and length upon Morn2 depletion, with restoration following rescue. Scale bar: 10 μm. (C’) Quantification of cilium length in E from three independent experiments with unpaired two-tailed t-tests is shown in F. Mean ± s.d. values are presented.

To determine whether gliding defects were due to deficient ciliogenesis, immunostaining for acetylated tubulin was performed. Morn2 depletion resulted in a significant reduction in motile cilia, which was restored by Morn2 MO-resistant mRNA (Figure 3B and 3B’). This phenotype was independently reproduced using a non-overlapping splice-blocking MO (sp-MO) (Figure S2A and S2A’), demonstrating that Morn2 is required for ciliogenesis of MCCs. Scanning electron microscopy (SEM) further confirmed that Morn2 knockdown led to a significant reduction in cilia number and length (Figure 3C and 3C’). Despite the profound impact on cilia formation, further analysis indicated that Morn2 knockdown did not alter basal body number or their apical docking (Figure S3A-S3A”). This observation aligns with its subcellular localization restricted to the axoneme and supports a model in which Morn2 acts downstream of basal body positioning to promote axoneme assembly and ciliary elongation.

### Ccdc30 Controls Centriole Amplification and MCC Development

Ccdc30 is a completely uncharacterized protein with no prior functional studies. Its striking ring-like localization surrounding the basal body prompted us to further investigate its function. Knockdown of Ccdc30 by a specific MO also severely impaired embryo gliding (Figure 4A and 4A’). Unlike Morn2 knockdown, Cilia staining showed that depletion of Ccdc30 not only impaired ciliogenesis but also decreased the overall population of MCCs (Figure 4B-4B’’). Similarly, Ccdc30 sp-MO produced similar ciliogenesis defects (Figure S2B and S2B’), indicating its involvement in both ciliogenesis and MCC development.

**Figure 4.**
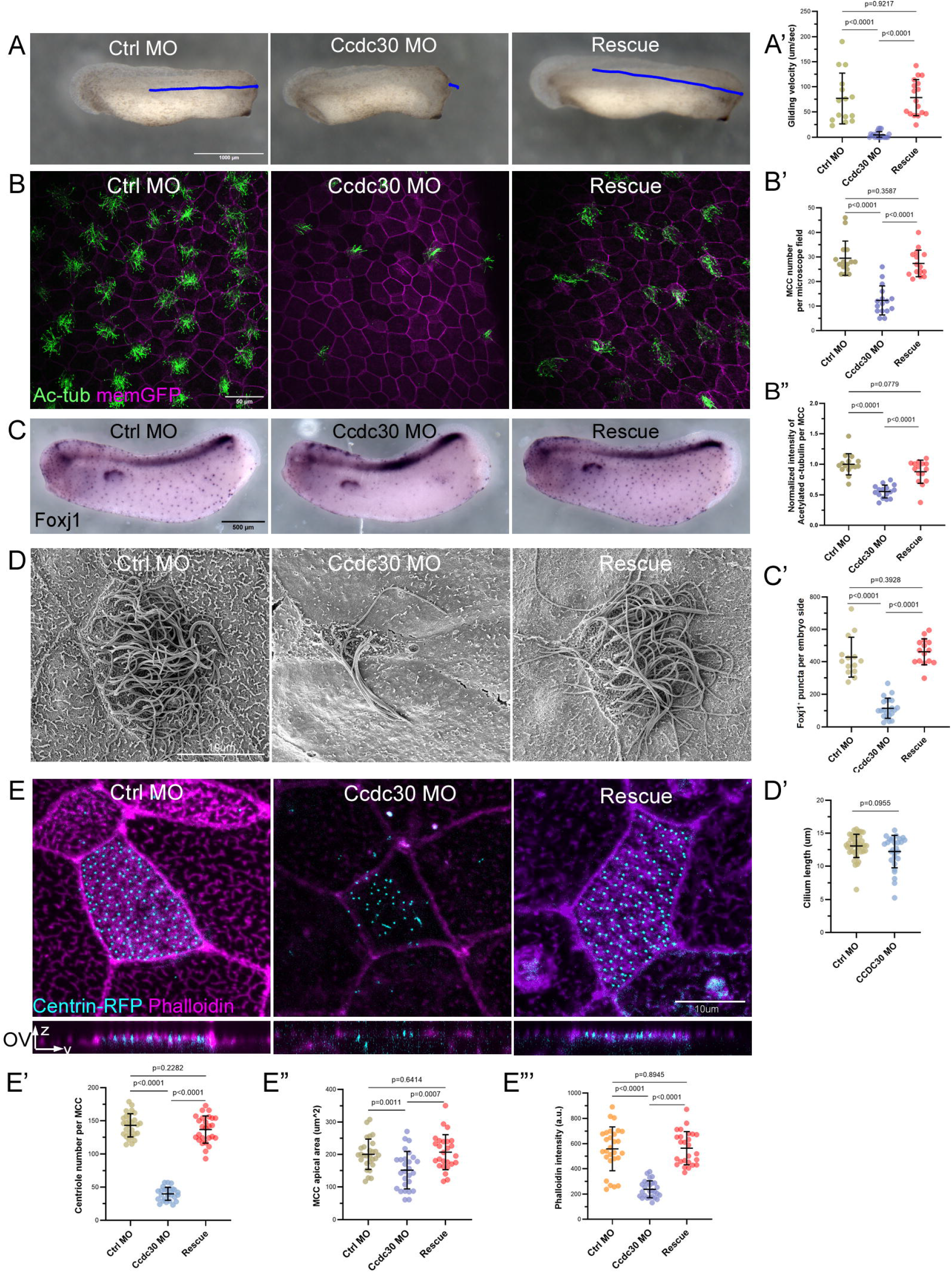
Ccdc30 regulates MCC mantainance, centriole amplification, and apical cytoskeletal organization. (A) Gliding assay showing that knockdown of Ccdc30 severely impaired embryo motility, which was rescued by co-injection of Ccdc30 MO-resistant mRNA. Scale bar: 1000 μm. (A’) Quantification of gliding velocity in A from three independent experiments with unpaired two-tailed t-tests is shown in B. Mean ± s.d. values are presented. (B) Immunostaining for acetylated tubulin demonstrates that Ccdc30 depletion reduces cilia number and MCC abundance. Scale bar: 50 μm. Quantification of normalized intensity of acetylated-tubulin (B’) and MCC number (B’’) in C from three independent experiments with unpaired two-tailed t-tests is shown in D and E, respectively. Mean ± s.d. values are presented. (C) Whole-mount in situ hybridization using the MCC marker *foxj1* confirms that Ccdc30 knockdown inhibits MCC formation, a phenotype reversed by Ccdc30 mRNA rescue. Scale bar: 500 μm. (C’) Quantification of MCC number in F from three independent experiments with unpaired two-tailed t-tests is shown in G. Mean ± s.d. values are presented. (D) Scanning electron microscopy reveals a dramatic reduction in cilia number and length in Ccdc30 morphants. Scale bar: 10 μm. (D’) Quantification of cilium length in H from three independent experiments with unpaired two-tailed t-tests is shown in I. Mean ± s.d. values are presented. (E) Phalloidin staining (actin) and Centrin4-RFP (basal bodies) show that in Ccdc30 morphants, centriole number is reduced, and basal body docking is impaired. Phalloidin signal at apical surface area is also significantly diminished. Scale bar: 10 μm. OV: orthogonal view. Quantification of centriole number (E’), apical surface area (E’’) and phalloidin intensity (E’’’) from three independent experiments with unpaired two-tailed t-tests, respectively. Mean ± s.d. values are presented.

Whole-mount in situ hybridization using the MCC specific marker *foxj1*, a transcriptional regulator of multiciliogenesis (Yu et al., 2008), further confirmed that depletion of Ccdc30 inhibited epidermal MCC abundance, a phenotype that could be rescued by co-injection of *ccdc30* MO-resistant mRNA (Figure 4C and 4C’). To examine whether Ccdc30 functions during MCC specification/induction, we examined *foxj1* expression at an earlier developmental stage during MCC specification (stage 14-15). At this stage, Ccdc30 depletion produced only a mild effect on the foxj1-positive MCC population (Figure S4A and S4A’) compared with the strong reduction observed at later stages (Figure 4C and 4C’). This result suggests that Ccdc30 is unlikely to be a regulator required for MCC specification/induction, but may contribute to the maintenance of the MCC population.

SEM analysis confirmed the reduction in cilia number without compromising cilia length, suggesting a possible function in regulating basal body abundance (Figure 4D and 4D’). Indeed, consistent with its centriole-associated localization, a marked reduction in centriole number was observed in Ccdc30 morphants (Figure 4E and 4E’), a defect that was fully rescued by co-injection of MO-resistant *ccdc30* mRNA, indicating a critical role for Ccdc30 in centriole amplification. Furthermore, considering that Ccdc30 forms a ring larger than that of the distal appendage marker Cep164, we examined whether Ccdc30 depletion alters Cep164 recruitment to centrioles. The accumulation of Cep164 was preserved in the majority of centrioles in Ccdc30 morphants (Figure S4B), indicating that distal appendage formation and centriole maturation are likely unaffected by Ccdc30 depletion.

Beyond centriole biogenesis, ciliogenesis also relies on the formation of the apical actin meshwork and the proper migration and docking of basal bodies to the apical surface (Zhao et al., 2022; Lee et al., 2022, 2019; Kulkarni et al., 2018; Antoniades et al., 2014; Park et al., 2006). In Ccdc30-depleted MCCs, basal bodies failed to dock efficiently, accompanied by a significant loss of phalloidin staining, which reflects disruption of the apical actin network. Consequently, the apical surface area of MCCs was significantly reduced (Figure 4E, 4E” and 4E”‘). Collectively, these findings suggest that, in contrast to Morn2, which functions directly in axoneme assembly, Ccdc30 plays broader roles during MCC development. At earlier stages, Ccdc30 may contribute to centriole amplification in MCCs; during ciliogenesis, it likely supports apical actin organization necessary for basal body migration and docking.

## DISCUSSION

Motile cilia are essential for fluid propulsion in diverse biological contexts, and their dysfunction contributes to a wide spectrum of human diseases (Brooks and Wallingford, 2014; Fliegauf et al., 2007; Spassky and Meunier, 2017). Despite extensive efforts to elucidate the molecular architecture of motile cilia, many regulators of ciliogenesis remain unknown, in part due to methodological limitations. Prior proteomic studies have predominantly focused on purified ciliary axonemes to enrich for ciliary proteins (Blackburn et al., 2017; Boesger et al., 2009; Broadhead et al., 2006; Ostrowski et al., 2002; Pazour et al., 2005; Sim et al., 2020; Subota et al., 2014; Li et al., 2004). This approach, while informative, excludes key regulatory components located in other cellular compartments such as basal bodies, cytoplasm, and nuclei. Our study addresses this gap by presenting a high-resolution, whole-cell proteomic profile of vertebrate MCCs, generated using *X. laevis* mucociliary organoids in combination with an MCIDAS-driven enrichment strategy and OFIC-based proteomics.

The use of a glucocorticoid-inducible MCIDAS expression system enabled robust MCC induction in animal cap organoids, creating a gain-of-function model that facilitated the enrichment of MCC-specific proteins. Comparing proteomes between Dex-treated and untreated samples allowed us to isolate proteins specifically associated with MCC identity and ciliogenesis. This strategy overcomes a key challenge in whole-cell proteomics—the presence of background signals from non-MCC cell types—and enhances the specificity of candidate identification. Through this approach, we identified over 9,000 proteins, with 832 significantly upregulated in MCCs, including well-established cilia-related proteins such as Ccp110, Cep164, and Dnai1. GO enrichment and database comparisons validated the quality and relevance of our dataset and highlighted its effectiveness in identifying novel regulators of ciliogenesis.

Beyond cataloging known ciliary components, this study is significant for uncovering previously uncharacterized proteins with distinct subcellular localizations and functions. For instance, phylogenetic analysis suggests Morn2 is an orthologue of *Chlamydomonas* C1a-18/Fap227 (Figure S3B), which localizes to the central apparatus (Wargo et al., 2005). The function of C1a-18 remains unknown, but our results are intriguing because while most central pair protein govern cilia beating (Samsel et al., 2021), some also control cilia assembly (Lechtreck et al., 2013), as we show here for *Xenopus* Morn2.

Ccdc30 was found to function at multiple levels of MCC development. It was required not only for centriole amplification but also for apical actin meshwork organization and basal body docking—processes that are essential for the formation of functional motile cilia. The localization of Ccdc30, which forms a larger ring-like structure distal to the known basal body appendage marker Cep164, suggests that it may define a novel subdomain of the basal body or associate with a distinct structural element involved in basal body maturation. One possibility is that Ccdc30 localizes to the base of the ciliary pocket, a region closely linked to actin organization. This could explain the disrupted apical actin meshwork observed in Ccdc30-depleted MCCs, consistent with the ciliary pocket’s known role in coordinating actin dynamics and vesicle trafficking (Benmerah, 2013).

Notably, Ccdc30 has been recently reported in association with asthma exacerbation in patients (Son et al., 2022). While this association does not directly implicate Ccdc30 in primary ciliary dyskinesia (PCD), a genetic disorder caused by defects in motile cilia structure and function, asthma is a well-recognized comorbidity in children with PCD (Zein et al., 2024). This clinical correlation further supports a functional link between Ccdc30 and motile cilia biology, raising the possibility that Ccdc30 plays a conserved role in the coordination of ciliary assembly and epithelial homeostasis relevant to human ciliopathies.

In conclusion, these findings demonstrate the power of whole-cell proteomics in combination with functional genomics to uncover new components and mechanisms of ciliogenesis. Importantly, the approach outlined here is generalizable and may be applied to other specialized cell types where proteome complexity and cell heterogeneity present analytical challenges. Furthermore, our data provide a rich resource for future studies of MCC development and cilia-related pathologies. This study not only expands the known repertoire of ciliary proteins but also sheds light on novel regulatory pathways in motile cilia formation and MCC differentiation. By moving beyond axoneme-centric methods, we reveal new layers of complexity in the assembly and function of motile cilia. These insights have broad implications for understanding the molecular basis of ciliopathies and for developing targeted therapeutic strategies to restore or enhance ciliary function in disease contexts.

## MATERIALS AND METHODS

### Animals

Wild-type *X. laevis* and *X. tropicalis* frogs were purchased from National *Xenopus* Resource in Woods Hole, Massachusetts. Embryos were obtained through in vitro fertilization and injected as described (Moody, 1999). Methods involving live animals were carried out in accordance with the guidelines and regulations approved and enforced by the Institutional Animal Care and Use Committees at University of Delaware.

### Plasmids and in vitro transcription

The expression construct for MCIDAS-GR was a courtesy of Brian Mitchell (Northwestern University). Coding sequences of *X. tropicalis* Ccdc30, Morn2, and Wdr88 were amplified by PCR from stage 18 and stage 25 embryo cDNA and subcloned into pCS2+-HA and pCS2+-GFP vectors. Capped sense RNAs were transcribed using the mMessage mMachine SP6 kit (Thermo Fisher).

### Microinjection of *X. laevis* embryos and animal cap dissection

One-cell stage *X. laevis* embryos were injected with 40 pg MCIDAS-GR mRNA. Animal caps were manually dissected from *X. laevis* embryos at stage 8 as described (Dingwell and Smith, 2018) and subsequently cultured in Danilchik’s for Amy medium supplemented with gentamycin (50 μg/ml). Protein activity of GR chimeric constructs was induced by incubating the embryos in 4 μg/ml dexamethasone from stage 11–26.

### Microinjection of *X. tropicalis* embryos

For *X. tropicalis* embryos, the following mRNAs were used for one-cell injection: memGFP (100 pg); Morn2-HA (250 pg); Ccdc30-HA (300 pg); Wdr88-GFP (200 pg); Morn2-GFP (200 pg); Ccdc30-GFP (250 pg); Centrin4-RFP (50 pg). The following mRNAs were used for injection into one ventral blastomere at four-cell stage: memGFP (50 pg); Morn2-HA (200 pg); Ccdc30-HA (200 pg).

The morpholinos were obtained from Gene Tools with the following sequences: standard control MO: 5′-CTAAACTTGTGGTTCTGGCGGATA-3′; Morn2 MO: 5’-TCCCTAGACAAGTTTCGTATCCATA-3’; Morn2 sp-MO: 5’-GTAATCATTTCTCACCACACGGTTC-3’; Ccdc30 MO: 5’-CATCCATTTTGGCTTCTTACTTCAG-3’; Ccdc30 sp-MO: 5’-AGAAGCAAAGCCATACCTCATCCAT-3’. 7 ng control MO, Morn2 MO or Ccdc30 MO was injected per *X. tropicalis* embryo at one-cell stage. 4 ng control MO, Morn2 sp-MO or Ccdc30 sp-MO was injected into one ventral blastomere at four-cell stage.

### Whole-mount in situ hybridization

Digoxigenin-labeled antisense RNA probes were synthesized using T3 RNA polymerase, and whole-mount in situ hybridization were performed for *X. tropicalis* embryos using probes for *foxj1, ccdc30, wdr88*, and *morn2* as described (Sive et al., 2010).

### Immunofluorescence

Embryos were fixed with 4% formaldehyde at 4°C overnight and then dehydrated with 100% methanol. The following primary antibodies were used: mouse anti-acetylated tubulin-594 (1:1000, Proteintech), rat anti-HA (1:500, Sigma) and rabbit anti-CEP164 (1:400, Proteintech). Secondary antibodies conjugated to Alexa Fluor dyes were used at 1:1000. The samples were mounted in ProLong Gold and imaged using an Andor Dragonfly spinning disk confocal microscope or Zeiss LSM880 laser scanning confocal microscope.

### Live imaging

For live imaging, embryos expressing GFP-tagged constructs were mounted in 0.1× MMR with. Z-stacks imaging was acquired with Andor Dragonfly spinning disk confocal microscope or Zeiss LSM700 laser scanning confocal microscope.

### Electron Microscopy

The embryos were fixed in 2% paraformaldehyde and 2% glutaraldehyde in 0.1M sodium cacodylate buffer, rinsed with the buffer, and then fixed in 1% osmium tetroxide in 0.1M sodium cacodylate buffer. They were subsequently dehydrated in ascending concentrations of ethanol (25% to 100%) followed by hexamethyldisilazane and air dried. The dried samples were mounted on SEM stubs and sputter coated with platinum in a Leica EM ACE600 coater (Vienna, Austria). SEM images were acquired using a ThermoFisher Scientific Apreo Volumescope SEM (Hillsboro, OR).

### Embryo gliding assay

Stage-27 embryos were placed in 0.1× MMR on agar-coated dishes. Time-lapse videos were captured using a Zeiss stereomicroscope. Movement trajectories were quantified using ImageJ’s Manual tracking plugin.

### Protein sample preparation and LC-MS/ analysis

For in-cell sample preparation, eight stage-26 *X. laevis* animal caps (Dex+ or Dex-) were collected and transferred to E4tips XL (CDS Analytical, Oxford, PA) that were prefilled with 200 µl of methanol and incubated on ice for 30 minutes. Three replicates were prepared. The tips containing fixed embryos were processed following a protocol described previously (Elsayyid et al., 2025; Sun et al., 2025). Briefly, the tips were first spun at 1,500 x g for one minute and washed one time with 200 µl of methanol. Then 100 µl of 50 mM triethylammonium bicarbonate (TEAB) with final concentration of 10 mM Tris (2-carboxyethyl) phosphine (TCEP) and 40 mM chloroacetamide (CAA) was added, followed by incubation at 45□C for 10-15 min. The E4tips were then washed one time with 200 µl 50 mM TEAB. For protein digestion, 150 µl 50 mM TEAB plus 1.0 µg of trypsin/Lys-C mix (Promega, WI) was added to the samples and incubated at 37□C for 16-18 hours with gentle shaking (350 rpm/min). After digestion, the samples were acidified with 1% formic acid (final concentration) and spun at low speed (400-600 x g) for 10 min. The tips were washed one time with 200 µl of wash buffer (0.5% acetic acid in water) to discard flow through. The tips were transferred to new collection tubes, and subjected to two sequential elution with 200 µl of Elution buffer I (60% acetonitrile and 0.5% acetic acid in water) and Elution buffer II (80% ACN and 0.5% acetic acid in water), respectively. The elution was pooled, dried in the SpeedVac, and then stored at −80□C until further analysis.

### LC-MS/MS analysis

The LC-MS/MS analysis was conducted with an Ultimate 3000 RSLCnano system coupled to an Orbitrap Eclipse mass spectrometer and FAIMS Pro Interface (Thermo Scientific). The peptides were resuspended in 15 µl of LC buffer A (0.1% formic acid in water), and 10 µl of it was loaded onto a trap column (PepMap100 C18, 300 μm × 2 mm, 5 μm; Thermo Scientific) followed by separation on an analytical column (PepMap100 C18, 50 cm × 75 μm i.d., 3 μm; Thermo Scientific) flowing at 250 nl/min. A linear LC gradient was applied from 1% to 25% mobile phase B over 125 min, followed by an increase to 32% mobile phase B over 10 min. The column was washed with 80% mobile phase B for 5 min, followed by equilibration with mobile phase A for 15 min. For the ion source settings, the spray voltage was set to 1.9 kV, funnel RF level at 50%, and heated capillary temperature at 275°C. The MS data were acquired in Orbitrap at 120K resolution, followed by MS/MS acquisition in data-independent mode following a protocol described previously (Martin et al., 2024). For FAIMS analysis, a 3-CV experiment (−40|-55|-75) was applied. The MS scan range (m/z) was set to 380-985, maximum injection time was 246ms, and normalized AGC target was 100%. For MS/MS acquisition, the isolation mode was Quadrupole, isolation window was 8 m/z, and window overlap was 1 m/z. The collision energy was 30%, Orbitrap resolution was 30K, AGC Target was 400K, and normalized AGC target was 800%.

### Proteome quantitation and data analysis

Mass spec data were processed using Spectronaut software (version 19.5) (Bruderer et al., 2015) and a library-free DIA analysis workflow with directDIA+ and the X. laevis protein database (UniProt 2025 release; 111,661 sequences). Briefly, the settings for Pulsar and library generation include: Trypsin/P as specific enzyme; peptide length from 7 to 52 amino acids; allowing 2 missed cleavages; toggle N-terminal M turned on; Carbamidomethyl on C as fixed modification; Oxidation on M and Acetyl at protein N-terminus as variable modifications; FDRs at PSM, peptide and protein level all set to 0.01; Quantity MS level set to MS2, and cross-run normalization turned on. Bioinformatics analyses including t-test, correlation, volcano plot and clustering analyses were performed using Perseus software (version 1.6.2.3) and Prism GraphPad (version 10) unless otherwise indicated. The MS raw files associated with this study have been deposited to the MassIVE server (https://massive.ucsd.edu/) with the dataset identifiers MSV000097879.

### Statistical Analysis

Statistical significance was assessed using unpaired two-tailed Student’s t-tests for comparisons between two groups. p-values <0.05 were considered statistically significant. All statistical analyses were performed using GraphPad Prism10.

## Data availability

Proteomic data have been deposited to the MassIVE server (https://massive.ucsd.edu/) with the dataset identifier MSV000097879 and are publicly available as of the date of publication.

## ACKNOWLEDGMENTS

This work is supported by NIH R01 DE029802 (to S.W.) and R01 HD085901 (to J.B.W.). Microscopy equipment was acquired with a shared instrumentation grant (S10 OD025165) and access was supported by NIH-NIGMS (P20 GM103446), NIGMS (P20 GM139760) and the State of Delaware. Access to the Orbitrap Eclipse MS instrument was supported by the Institutional Development Award (IDeA) from the National Institute of Health’s National Institute of General Medical Sciences under grant number P20GM103446 and P20 GM104316. The MCIDAS-GR construct was kindly provided by Brian Mitchell. The table of content figure and some of the main figures were created with BioRender.

## AUTHOR CONTRIBUTIONS

J.S. conceived and designed the study. J.S. and X.X. performed the majority of the experiments, with assistance from T.Z. and F.C.. J.R. conducted scanning electron microscopy. Y.Y. carried out the LC-MS. A.L.P.T. provided *X. laevis* animals. J.S. drafted the manuscript. X.X. and N.S. analyzed the data and prepared the figures. S.W. supervised the project and revised the manuscript. J.W. offered guidance during revision and reviewed the manuscript. All authors contributed to data interpretation and approved the final version.

## SUPPLEMENTAL INFORMATION

**Document S1**. Source file for FigureS1-S4

**Video S1. Immunostaining of Ccdc30-HA and Cep164 in *X. tropicalis* epidermal MCC**. Centrin4-RFP (cyan), Cep164 (magenta), and Ccdc30-HA (green) are shown. The image that was cropped from Figure 2E highlights two basal bodies to visualize subcellular localization in greater detail. The video includes rotational views of the basal bodies to illustrate the spatial relationship between Ccdc30 and Cep164, a known distal appendage marker.

**Video S2. 3D view of live imaging of Ccdc30-GFP and Cep164-RFP in epidermal MCC**. Centrin4-BFP (magenta), Cep164-RFP (red), and Ccdc30-GFP (green) are shown.

**Table S1. MCC Proteome identified in this study. Summary of protein and peptide IDs**. List of all the identified proteins exported from Spectronaut, with description information. Pair-wise comparison of MCIDAS-GR +Dex and −Dex.

**Table S2. Human orthologs of *X. laevis* MCC-enriched proteins identified in this study**.

**Table S3. Overlap between proteins from this study and those in CiliaCarta**. Listed are the CiliaCarta dataset, genes found in common between these two datasets and genes unique in our dataset.

**Table S4. Overlap between proteins from this study and MCC core dataset**. Listed are *X. tropicalis* orthologs of the MCC-enriched proteins identified in this study, MCC core dataset, proteins found in common between these two datasets, and proteins unique in our dataset.

**Table S5. Overlap between proteins from this study and MCC scRNA-seq dataset**.

**Table S6. Overlap between proteins from this study and axoneme components**. Listed are axoneme components, proteins found in common between these two datasets, and proteins unique in our dataset.

**Table S7. Overlap between proteins from this study and Rfx2 direct targets**. Listed are Rfx2 direct target dataset, genes found in common between these two datasets, and genes unique in our dataset.

## REFERENCES

Angerilli, A., P. Smialowski, and R.A. Rupp. 2018. The Xenopus animal cap transcriptome: building a mucociliary epithelium. Nucleic Acids Research. 46:8772–8787. doi:10.1093/nar/gky771.

Antoniades, I., P. Stylianou, and P.A. Skourides. 2014. Making the Connection: Ciliary Adhesion Complexes Anchor Basal Bodies to the Actin Cytoskeleton. Developmental Cell. 28:70–80. doi:10.1016/j.devcel.2013.12.003.

Benmerah, A. 2013. The ciliary pocket. Current Opinion in Cell Biology. 25:78–84. doi:10.1016/j.ceb.2012.10.011.

Blackburn, K., X. Bustamante-Marin, W. Yin, M.B. Goshe, and L.E. Ostrowski. 2017. Quantitative Proteomic Analysis of Human Airway Cilia Identifies Previously Uncharacterized Proteins of High Abundance. J. Proteome Res. 16:1579–1592. doi:10.1021/acs.jproteome.6b00972.

Boesger, J., V. Wagner, W. Weisheit, and M. Mittag. 2009. Analysis of Flagellar Phosphoproteins from Chlamydomonas reinhardtii. Eukaryot Cell. 8:922–932. doi:10.1128/EC.00067-09.

Boutin, C., and L. Kodjabachian. 2019. Biology of multiciliated cells. Current Opinion in Genetics & Development. 56:1–7. doi:10.1016/j.gde.2019.04.006.

Broadhead, R., H.R. Dawe, H. Farr, S. Griffiths, S.R. Hart, N. Portman, M.K. Shaw, M.L. Ginger, S.J. Gaskell, P.G. McKean, and K. Gull. 2006. Flagellar motility is required for the viability of the bloodstream trypanosome. Nature. 440:224–227. doi:10.1038/nature04541.

Brooks, E.R., and J.B. Wallingford. 2014. Multiciliated Cells. Current Biology. 24:R973–R982. doi:10.1016/j.cub.2014.08.047.

Bruderer, R., O.M. Bernhardt, T. Gandhi, S.M. Miladinović, L.-Y. Cheng, S. Messner, T. Ehrenberger, V. Zanotelli, Y. Butscheid, C. Escher, O. Vitek, O. Rinner, and L. Reiter. 2015. Extending the Limits of Quantitative Proteome Profiling with Data-Independent Acquisition and Application to Acetaminophen-Treated Three-Dimensional Liver Microtissues. Molecular & Cellular Proteomics. 14:1400–1410. doi:10.1074/mcp.M114.044305.

Chen, X., Z. Shi, F. Yang, T. Zhou, and S. Xie. 2023. Deciphering cilia and ciliopathies using proteomic approaches. The FEBS Journal. 290:2590–2603. doi:10.1111/febs.16538.

Chung, M.-I., T. Kwon, F. Tu, E.R. Brooks, R. Gupta, M. Meyer, J.C. Baker, E.M. Marcotte, and J.B. Wallingford. 2014. Coordinated genomic control of ciliogenesis and cell movement by RFX2. eLife. 3:e01439. doi:10.7554/eLife.01439.

Dingwell, K.S., and J.C. Smith. 2018. Dissecting and Culturing Animal Cap Explants. Cold Spring Harb Protoc. 2018:pdb.prot097329. doi:10.1101/pdb.prot097329.

Elsayyid, M., J.E. Tanis, and Y. Yu. 2025. Simple In-Cell Processing Enables Deep Proteome Analysis of Low-Input Caenorhabditis elegans. Anal. Chem. 97:9159–9167. doi:10.1021/acs.analchem.4c05003.

Fliegauf, M., T. Benzing, and H. Omran. 2007. When cilia go bad: cilia defects and ciliopathies. Nat Rev Mol Cell Biol. 8:880–893. doi:10.1038/nrm2278.

Guo, T., C. Lu, D. Yang, C. Lei, Y. Liu, Y. Xu, B. Yang, R. Wang, and H. Luo. 2022. Case Report: DNAAF4 Variants Cause Primary Ciliary Dyskinesia and Infertility in Two Han Chinese Families. Front. Genet. 13:934920. doi:10.3389/fgene.2022.934920.

Hellsten, U., R.M. Harland, M.J. Gilchrist, D. Hendrix, J. Jurka, V. Kapitonov, I. Ovcharenko, N.H. Putnam, S. Shu, L. Taher, I.L. Blitz, B. Blumberg, D.S. Dichmann, I. Dubchak, E. Amaya, J.C. Detter, R. Fletcher, D.S. Gerhard, D. Goodstein, T. Graves, I.V. Grigoriev, J. Grimwood, T. Kawashima, E. Lindquist, S.M. Lucas, P.E. Mead, T. Mitros, H. Ogino, Y. Ohta, A.V. Poliakov, N. Pollet, J. Robert, A. Salamov, A.K. Sater, J. Schmutz, A. Terry, P.D. Vize, W.C. Warren, D. Wells, A. Wills, R.K. Wilson, L.B. Zimmerman, A.M. Zorn, R. Grainger, T. Grammer, M.K. Khokha, P.M. Richardson, and D.S. Rokhsar. 2010. The Genome of the Western Clawed Frog Xenopus tropicalis. Science. 328:633–636. doi:10.1126/science.1183670.

Kott, E., M. Legendre, B. Copin, J.-F. Papon, F. Dastot-Le Moal, G. Montantin, P. Duquesnoy, W. Piterboth, D. Amram, L. Bassinet, J. Beucher, N. Beydon, E. Deneuville, V. Houdouin, H. Journel, J. Just, N. Nathan, A. Tamalet, N. Collot, L. Jeanson, M. Le Gouez, B. Vallette, A.-M. Vojtek, R. Epaud, A. Coste, A. Clement, B. Housset, B. Louis, E. Escudier, and S. Amselem. 2013. Loss-of-Function Mutations in RSPH1 Cause Primary Ciliary Dyskinesia with Central-Complex and Radial-Spoke Defects. The American Journal of Human Genetics. 93:561–570. doi:10.1016/j.ajhg.2013.07.013.

Kulkarni, S.S., J.N. Griffin, P.P. Date, K.F. Liem, and M.K. Khokha. 2018. WDR5 Stabilizes Actin Architecture to Promote Multiciliated Cell Formation. Developmental Cell. 46:595-610.e3. doi:10.1016/j.devcel.2018.08.009.

Lechtreck, K.-F., T.J. Gould, and G.B. Witman. 2013. Flagellar central pair assembly in Chlamydomonas reinhardtii. Cilia. 2:15. doi:10.1186/2046-2530-2-15.

Lee, J., A.F. Møller, S. Chae, A. Bussek, T.J. Park, Y. Kim, H.-S. Lee, T.H. Pers, T. Kwon, J. Sedzinski, and K.N. Natarajan. 2023. A single-cell, time-resolved profiling of Xenopus mucociliary epithelium reveals nonhierarchical model of development. Sci. Adv. 9:eadd5745. doi:10.1126/sciadv.add5745.

Lee, M., Y.-S. Hwang, J. Yoon, J. Sun, A. Harned, K. Nagashima, and I.O. Daar. 2019. Developmentally regulated GTP-binding protein 1 modulates ciliogenesis via an interaction with Dishevelled. Journal of Cell Biology. 218:2659–2676. doi:10.1083/jcb.201811147.

Lee, M., K. Nagashima, J. Yoon, J. Sun, Z. Wang, C. Carpenter, H.-K. Lee, Y.-S. Hwang, C.J. Westlake, and I.O. Daar. 2022. CEP97 phosphorylation by Dyrk1a is critical for centriole separation during multiciliogenesis. Journal of Cell Biology. 221:e202102110. doi:10.1083/jcb.202102110.

Leung, M.R., C. Sun, J. Zeng, J.R. Anderson, Q. Niu, W. Huang, W.E.M. Noteborn, A. Brown, T. Zeev-Ben-Mordehai, and R. Zhang. 2025a. Structural diversity of axonemes across mammalian motile cilia. Nature. 637:1170–1177. doi:10.1038/s41586-024-08337-5.

Leung, M.R., C. Sun, J. Zeng, J.R. Anderson, Q. Niu, W. Huang, W.E.M. Noteborn, A. Brown, T. Zeev-Ben-Mordehai, and R. Zhang. 2025b. Structural diversity of axonemes across mammalian motile cilia. Nature. 637:1170–1177. doi:10.1038/s41586-024-08337-5.

Li, J.B., J.M. Gerdes, C.J. Haycraft, Y. Fan, T.M. Teslovich, H. May-Simera, H. Li, O.E. Blacque, L. Li, C.C. Leitch, R.A. Lewis, J.S. Green, P.S. Parfrey, M.R. Leroux, W.S. Davidson, P.L. Beales, L.M. Guay-Woodford, B.K. Yoder, G.D. Stormo, N. Katsanis, and S.K. Dutcher. 2004. Comparative Genomics Identifies a Flagellar and Basal Body Proteome that Includes the BBS5 Human Disease Gene. Cell. 117:541–552. doi:10.1016/S0092-8674(04)00450-7.

Ma, L., I. Quigley, H. Omran, and C. Kintner. 2014. Multicilin drives centriole biogenesis via E2f proteins. Genes Dev. 28:1461–1471. doi:10.1101/gad.243832.114.

Martin, K.R., H.T. Le, A. Abdelgawad, C. Yang, G. Lu, J.L. Keffer, X. Zhang, Z. Zhuang, P.N. Asare-Okai, C.S. Chan, M. Batish, and Y. Yu. 2024. Development of an efficient, effective, and economical technology for proteome analysis. Cell Reports Methods. 4:100796. doi:10.1016/j.crmeth.2024.100796.

Moody, S.A. 1999. Cell Lineage Analysis in Xenopus Embryos. In Developmental Biology Protocols. Humana Press, New Jersey. 331–347.

Nigg, E.A., and J.W. Raff. 2009. Centrioles, Centrosomes, and Cilia in Health and Disease. Cell. 139:663–678. doi:10.1016/j.cell.2009.10.036.

Onoufriadis, A., A. Shoemark, M. Schmidts, M. Patel, G. Jimenez, H. Liu, B. Thomas, M. Dixon, R.A. Hirst, A. Rutman, T. Burgoyne, C. Williams, J. Scully, F. Bolard, J.-J. Lafitte, P.L. Beales, C. Hogg, P. Yang, E.M.K. Chung, R.D. Emes, C. O’Callaghan, UK10K, P. Bouvagnet, and H.M. Mitchison. 2014. Targeted NGS gene panel identifies mutations in RSPH1 causing primary ciliary dyskinesia and a common mechanism for ciliary central pair agenesis due to radial spoke defects. Human Molecular Genetics. 23:3362–3374. doi:10.1093/hmg/ddu046.

Ostrowski, L.E., K. Blackburn, K.M. Radde, M.B. Moyer, D.M. Schlatzer, A. Moseley, and R.C. Boucher. 2002. A Proteomic Analysis of Human Cilia. Molecular & Cellular Proteomics. 1:451–465. doi:10.1074/mcp.M200037-MCP200.

Park, T.J., S.L. Haigo, and J.B. Wallingford. 2006. Ciliogenesis defects in embryos lacking inturned or fuzzy function are associated with failure of planar cell polarity and Hedgehog signaling. Nat Genet. 38:303–311. doi:10.1038/ng1753.

Park, T.J., B.J. Mitchell, P.B. Abitua, C. Kintner, and J.B. Wallingford. 2008. Dishevelled controls apical docking and planar polarization of basal bodies in ciliated epithelial cells. Nat Genet. 40:871–879. doi:10.1038/ng.104.

Pazour, G.J., N. Agrin, J. Leszyk, and G.B. Witman. 2005. Proteomic analysis of a eukaryotic cilium. The Journal of Cell Biology. 170:103–113. doi:10.1083/jcb.200504008.

Quigley, I.K., and C. Kintner. 2017. Rfx2 Stabilizes Foxj1 Binding at Chromatin Loops to Enable Multiciliated Cell Gene Expression. PLoS Genet. 13:e1006538. doi:10.1371/journal.pgen.1006538.

Reiter, J.F., and M.R. Leroux. 2017. Genes and molecular pathways underpinning ciliopathies. Nat Rev Mol Cell Biol. 18:533–547. doi:10.1038/nrm.2017.60.

Samsel, Z., J. Sekretarska, A. Osinka, D. Wloga, and E. Joachimiak. 2021. Central Apparatus, the Molecular Kickstarter of Ciliary and Flagellar Nanomachines. IJMS. 22:3013. doi:10.3390/ijms22063013.

Schmidt, K.N., S. Kuhns, A. Neuner, B. Hub, H. Zentgraf, and G. Pereira. 2012. Cep164 mediates vesicular docking to the mother centriole during early steps of ciliogenesis. Journal of Cell Biology. 199:1083–1101. doi:10.1083/jcb.201202126.

Session, A.M., Y. Uno, T. Kwon, J.A. Chapman, A. Toyoda, S. Takahashi, A. Fukui, A. Hikosaka, A. Suzuki, M. Kondo, S.J. Van Heeringen, I. Quigley, S. Heinz, H. Ogino, H. Ochi, U. Hellsten, J.B. Lyons, O. Simakov, N. Putnam, J. Stites, Y. Kuroki, T. Tanaka, T. Michiue, M. Watanabe, O. Bogdanovic, R. Lister, G. Georgiou, S.S. Paranjpe, I. Van Kruijsbergen, S. Shu, J. Carlson, T. Kinoshita, Y. Ohta, S. Mawaribuchi, J. Jenkins, J. Grimwood, J. Schmutz, T. Mitros, S.V. Mozaffari, Y. Suzuki, Y. Haramoto, T.S. Yamamoto, C. Takagi, R. Heald, K. Miller, C. Haudenschild, J. Kitzman, T. Nakayama, Y. Izutsu, J. Robert, J. Fortriede, K. Burns, V. Lotay, K. Karimi, Y. Yasuoka, D.S. Dichmann, M.F. Flajnik, D.W. Houston, J. Shendure, L. DuPasquier, P.D. Vize, A.M. Zorn, M. Ito, E.M. Marcotte, J.B. Wallingford, Y. Ito, M. Asashima, N. Ueno, Y. Matsuda, G.J.C. Veenstra, A. Fujiyama, R.M. Harland, M. Taira, and D.S. Rokhsar. 2016. Genome evolution in the allotetraploid frog Xenopus laevis. Nature. 538:336–343. doi:10.1038/nature19840.

Sim, H.J., S. Yun, H.E. Kim, K.Y. Kwon, G.-H. Kim, S. Yun, B.G. Kim, K. Myung, T.J. Park, and T. Kwon. 2020. Simple Method To Characterize the Ciliary Proteome of Multiciliated Cells. J. Proteome Res. 19:391–400. doi:10.1021/acs.jproteome.9b00589.

Sive, H.L., R. Grainger, and R.M. Harland. 2010. Early development of Xenopus laevis: a laboratory manual. Cold Spring Harbor Laboratory Press., Cold Spring Harbor, N.Y.

Son, J.-H., J.-S. Park, J.-U. Lee, M.K. Kim, S.-A. Min, C.-S. Park, and H.S. Chang. 2022. A genome-wide association study on frequent exacerbation of asthma depending on smoking status. Respiratory Medicine. 199:106877. doi:10.1016/j.rmed.2022.106877.

Spassky, N., and A. Meunier. 2017. The development and functions of multiciliated epithelia. Nat Rev Mol Cell Biol. 18:423–436. doi:10.1038/nrm.2017.21.

Spektor, A., W.Y. Tsang, D. Khoo, and B.D. Dynlacht. 2007. Cep97 and CP110 Suppress a Cilia Assembly Program. Cell. 130:678–690. doi:10.1016/j.cell.2007.06.027.

Stubbs, J.L., E.K. Vladar, J.D. Axelrod, and C. Kintner. 2012. Multicilin promotes centriole assembly and ciliogenesis during multiciliate cell differentiation. Nat Cell Biol. 14:140–147. doi:10.1038/ncb2406.

Subota, I., D. Julkowska, L. Vincensini, N. Reeg, J. Buisson, T. Blisnick, D. Huet, S. Perrot, J. Santi-Rocca, M. Duchateau, V. Hourdel, J.-C. Rousselle, N. Cayet, A. Namane, J. Chamot-Rooke, and P. Bastin. 2014. Proteomic Analysis of Intact Flagella of Procyclic Trypanosoma brucei Cells Identifies Novel Flagellar Proteins with Unique Sub-localization and Dynamics. Molecular & Cellular Proteomics. 13:1769–1786. doi:10.1074/mcp.M113.033357.

Sun, J., X. Xu, S. Wei, and Y. Yu. 2025. In-cell Proteomics Enables High-Resolution Spatial and Temporal Mapping of Early Xenopus tropicalis Embryos. Molecular & Cellular Proteomics. 101481. doi:10.1016/j.mcpro.2025.101481.

Van Dam, T.J.P., J. Kennedy, R. Van Der Lee, E. De Vrieze, K.A. Wunderlich, S. Rix, G.W. Dougherty, N.J. Lambacher, C. Li, V.L. Jensen, M.R. Leroux, R. Hjeij, N. Horn, Y. Texier, Y. Wissinger, J. Van Reeuwijk, G. Wheway, B. Knapp, J.F. Scheel, B. Franco, D.A. Mans, E. Van Wijk, F. Képès, G.G. Slaats, G. Toedt, H. Kremer, H. Omran, K. Szymanska, K. Koutroumpas, M. Ueffing, T.-M.T. Nguyen, S.J.F. Letteboer, M.M. Oud, S.E.C. Van Beersum, M. Schmidts, P.L. Beales, Q. Lu, R.H. Giles, R. Szklarczyk, R.B. Russell, T.J. Gibson, C.A. Johnson, O.E. Blacque, U. Wolfrum, K. Boldt, R. Roepman, V. Hernandez-Hernandez, and M.A. Huynen. 2019. CiliaCarta: An integrated and validated compendium of ciliary genes. PLoS ONE. 14:e0216705. doi:10.1371/journal.pone.0216705.

Walentek, P. 2021. Xenopus epidermal and endodermal epithelia as models for mucociliary epithelial evolution, disease, and metaplasia. Genesis. 59:e23406. doi:10.1002/dvg.23406.

Walentek, P., and I.K. Quigley. 2017. What we can learn from a tadpole about ciliopathies and airway diseases: Using systems biology in Xenopus to study cilia and mucociliary epithelia. Genesis. 55:e23001. doi:10.1002/dvg.23001.

Wargo, M.J., E.E. Dymek, and E.F. Smith. 2005. Calmodulin and PF6 are components of a complex that localizes to the C1 microtubule of the flagellar central apparatus. Journal of Cell Science. 118:4655–4665. doi:10.1242/jcs.02585.

Werner, M.E., and B.J. Mitchell. 2012. Understanding ciliated epithelia: The power of Xenopus. Genesis. 50:176–185. doi:10.1002/dvg.20824.

Werner, M.E., and B.J. Mitchell. 2013. Using Xenopus Skin to Study Cilia Development and Function. In Methods in Enzymology. Elsevier. 191–217.

Yu, X., C.P. Ng, H. Habacher, and S. Roy. 2008. Foxj1 transcription factors are master regulators of the motile ciliogenic program. Nat Genet. 40:1445–1453. doi:10.1038/ng.263.

Zariwala, M.A., M.W. Leigh, F. Ceppa, M.P. Kennedy, P.G. Noone, J.L. Carson, M.J. Hazucha, A. Lori, J. Horvath, H. Olbrich, N.T. Loges, A.-M. Bridoux, G. Pennarun, B. Duriez, E. Escudier, H.M. Mitchison, R. Chodhari, E.M.K. Chung, L.C. Morgan, R.U. De Iongh, J. Rutland, U. Pradal, H. Omran, S. Amselem, and M.R. Knowles. 2006. Mutations of DNAI1 in Primary Ciliary Dyskinesia: Evidence of Founder Effect in a Common Mutation. Am J Respir Crit Care Med. 174:858–866. doi:10.1164/rccm.200603-370OC.

Zein, J., A. Owora, H.J. Kim, N. Marozkina, and B. Gaston. 2024. Asthma Among Children With Primary Ciliary Dyskinesia. JAMA Netw Open. 7:e2449795. doi:10.1001/jamanetworkopen.2024.49795.

Zhao, H., J. Sun, C. Insinna, Q. Lu, Z. Wang, K. Nagashima, J. Stauffer, T. Andresson, S. Specht, S. Perera, I.O. Daar, and C.J. Westlake. 2022. Male infertility-associated Ccdc108 regulates multiciliogenesis via the intraflagellar transport machinery. EMBO Reports. 23:e52775. doi:10.15252/embr.202152775.

