## Supplemental Figure S1-S4 for "Whole-Cell Proteomics Identifies Novel Regulators of Ciliogenesis Beyond the Axoneme"

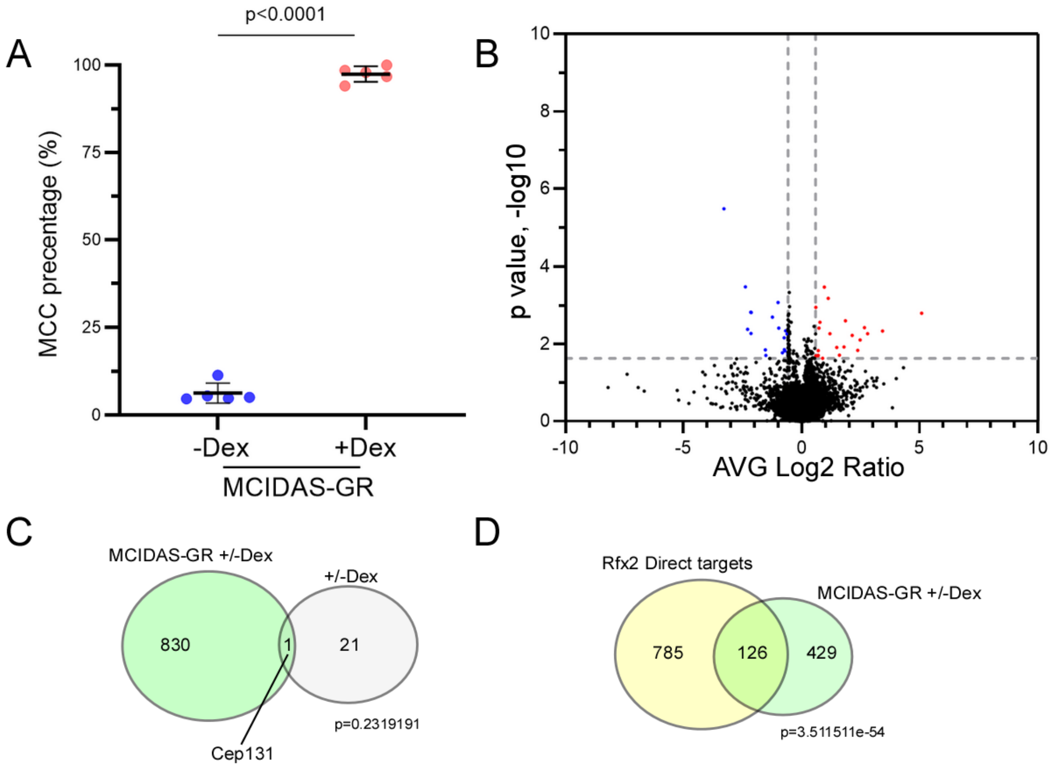


**Figure S1. Quantification of MCCs population and dataset comparisions**

(A) Quantification of MCCs from MCIDAS-GR injected animal cap organoids with or without Dex treatment.

(B) Volcano plot comparing Dex-treated and untreated organoids.

(C) The accompanying Venn diagram shows the overlap of upregulated proteins between the Dex-only comparison (±Dex) and the MCC-enriched condition (MCIDAS-hGR ±Dex).

(D) Venn diagram showing the overlap between Rfx2 direct target genes and our MCC proteome dataset.

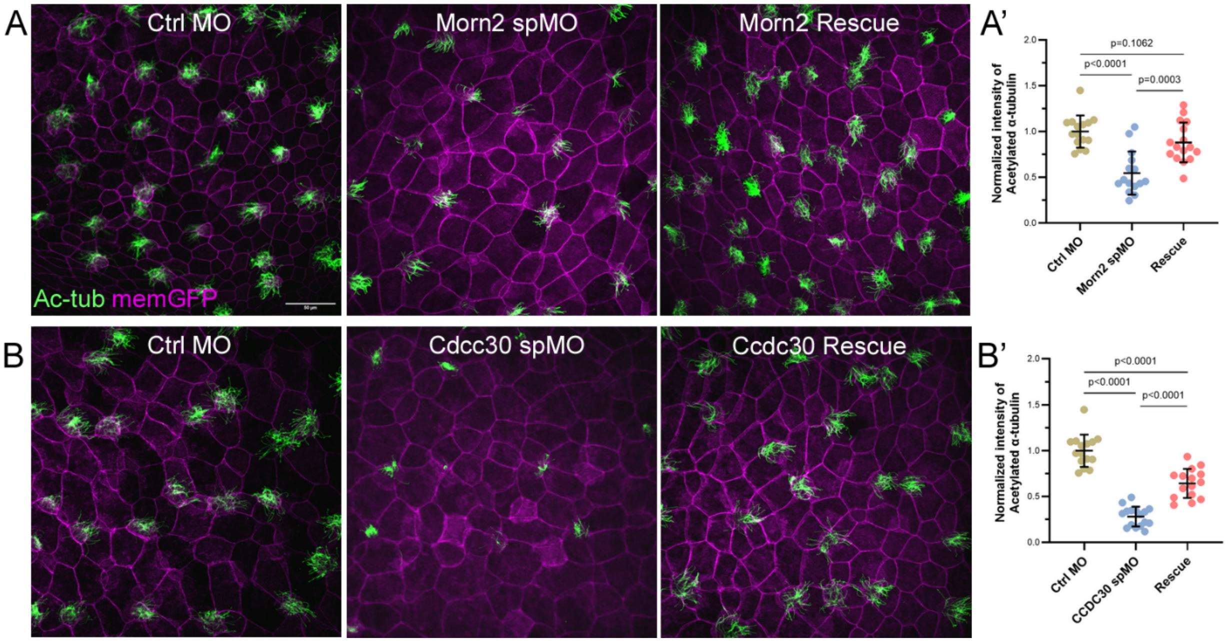


**Figure S2. Knockdown of *morn2* or *ccdc30* using splice-blocking morpholinos impairs ciliogenesis.**

(A,B) Immunostaining for acetylated tubulin in epidermal MCCs shows reduced ciliation following splice-blocking morpholino (sp-MO) knockdown of morn2 (A) or ccdc30 (B). Scale bars, 50 µm.

(A',B') Quantification of normalized acetylated-tubulin signal intensity from (A,B) across three independent experiments. Data are presented as mean ± s.d.; statistical significance was assessed using unpaired two-tailed t-tests.

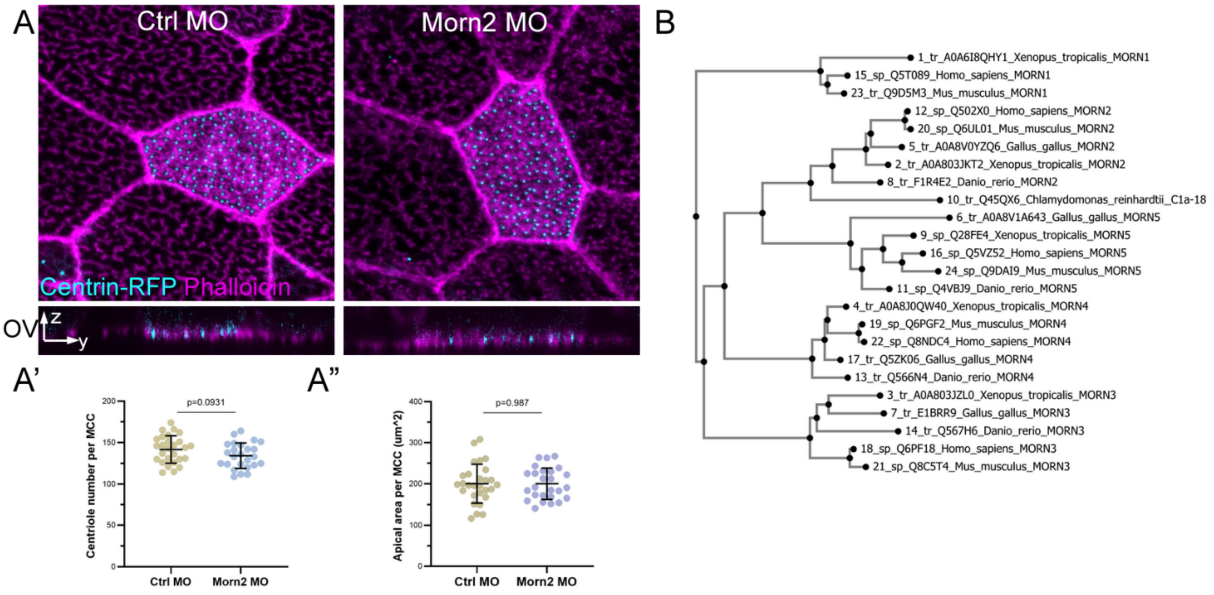


**Figure S3. Morn2 is dispensable for centriole amplification or apical docking.** (A) Phalloidin staining for actin network and Centrin4-RFP for basal bodies indicate that basal body number and apical docking remain largely unaffected in Morn2 morphants. Scale bar: 10 µm. (A',A") Quantification of centriole number per MCC and apical surface area in G from three independent experiments with unpaired two-tailed t-tests is shown in H and I, respectively. Mean ± s.d. values are presented. ns: no statistical differences between the groups.

(B) Phylogenetic tree based on Morn proteins and C1a-18 cross species. Full-length amino-acid sequence alignment and the neighbor joining tree was created with MAFFT, with the tree visualized using Phylo.io.

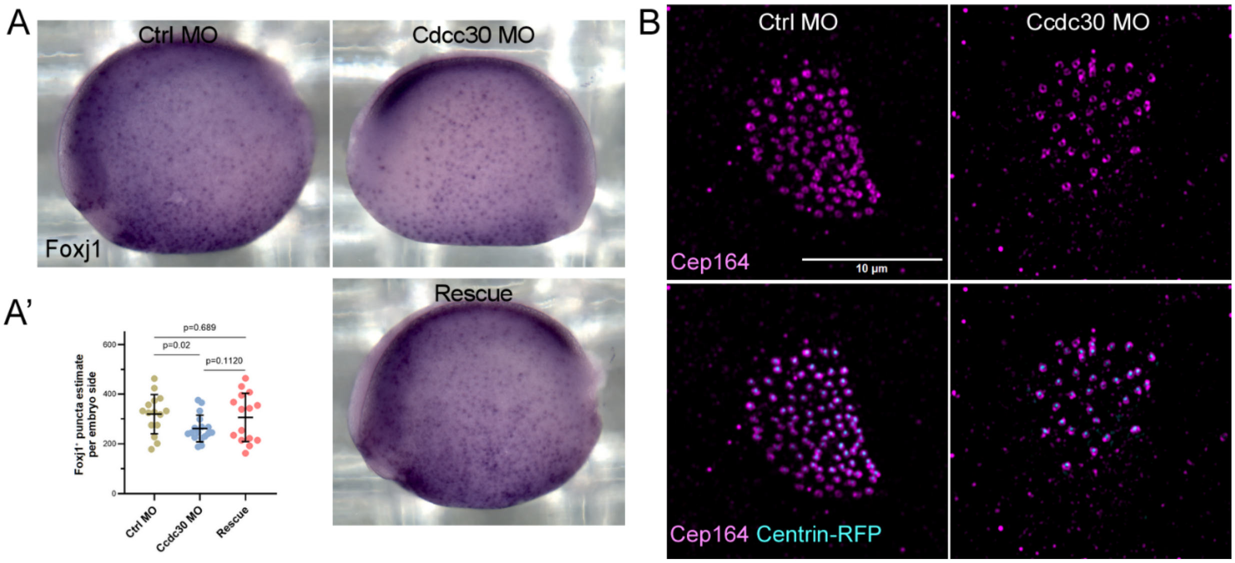


**Figure S4. Ccdc30 depletion does not markedly disrupt early MCC specification and reveals conserved ring-like localization.**

(A) Whole-mount in situ hybridization for the MCC marker foxj1 shows that Ccdc30 knockdown has minimal effect on early MCC induction/specification. MCC number was quantified from three independent experiments. Data are mean ± s.d.; statistical significance was assessed using unpaired two-tailed *t*-tests.

(B) Immunostaining for Cep164 in control and Ccdc30 morphants indicates that Ccdc30 depletion does not impair Cep164 recruitment to basal body distal appendages.
